# Impact of Different Acoustic Components on EEG-based Auditory Attention Decoding in Noisy and Reverberant Conditions

**DOI:** 10.1101/360834

**Authors:** Ali Aroudi, Bojana Mirkovic, Maarten De Vos, Simon Doclo

## Abstract

Recently, a least-squares-based method has been proposed to decode auditory attention from single-trial EEG recordings for an acoustic scenario with two competing speakers. This method aims at reconstructing the attended speech envelope from the EEG recordings using a trained spatio-temporal filter. While the performance of this method has been mainly studied for noiseless and anechoic acoustic conditions, it is important to fully understand its performance in realistic noisy and reverberant acoustic conditions. In this paper, we investigate auditory attention decoding (AAD) using EEG recordings for different acoustic conditions (anechoic, reverberant, noisy, and reverberant-noisy). In particular, we investigate the impact of different acoustic conditions for AAD filter training and for decoding. In addition, we investigate the influence on the decoding performance of the different acoustic components (i.e. reverberation, background noise and interfering speaker) in the reference signals used for decoding and the training signals used for computing the filters. First, we found that for all considered acoustic conditions it is possible to decode auditory attention with a decoding performance larger than 90%, even when the acoustic conditions for AAD filter training and for decoding are different. Second, when using reference signals affected by reverberation and/or background noise, a comparable decoding performance as when using clean reference signals can be obtained. In contrast, when using reference signals affected by the interfering speaker, the decoding performance significantly decreases. Third, the experimental results indicate that it is even feasible to use training signals affected by reverberation, background noise and/or the interfering speaker for computing the filters.

## I. INTRODUCTION

In complex acoustic conditions the human auditory system has a remarkable ability to segregate a speaker of interest from a mixture of speakers and background noise [1], [2]. In contrast with normal-hearing persons, hearing-impaired persons typically have more difficulties with such auditory segregation, particularly in multi-talker scenarios [3]. Although many acoustic signal processing algorithms are available to reduce background noise or to perform source separation in multi-talker scenarios [4], [5], these algorithms typically need to rely on assumptions about the target speaker to be enhanced. For example, in hearing aid applications the target speaker is typically assumed to be located in front of the user or is assumed to be the loudest speaker. As in real-world conditions such assumptions are often violated, the performance of these algorithms may substantially decrease. Therefore, successfully identifying the target speaker in hearing aid applications is very important to improve speech intelligibility.

Recent studies have shown that auditory cortical responses are correlated with the envelope of the attended speech signal [6]–[9]. Based on this finding, an auditory attention decoding (AAD) method has been proposed in [10] to identify the attended speaker from single-trial EEG recordings. This method aims at reconstructing the attended speech envelope from the EEG recordings using a trained spatio-temporal filter. In the *training step*, the clean speech signal of the attended speaker is used to train a spatio-temporal filter by minimizing the least-squares error between the attended speech envelope and the reconstructed envelope. In the *decoding step*, the clean speech signals of both the attended and the unattended speaker are used as reference signals. In [10] it has been shown that for high-density EEG recordings it is possible to decode auditory attention when presenting the clean speech signals of the different speakers to different ears of a listener (i.e. dichotic stimuli presentation). When presenting competing speech signals in a simulated anechoic condition including head filtering effects, it has been shown in [11] that a larger AAD performance can be obtained compared to dichotic presentation. Recently, a large research effort has focused on investigating how to use AAD as part of a brain-computer interface for real-world applications, e.g., to control a hearing aid [11]–[22], mainly however for anechoic conditions. Aiming at integrating a small-size EEG recording system in hearing aids, in [12]–[14] the reliability of AAD using a low number of EEG electrodes has been shown in an anechoic condition. Aiming at investigating the effect of neurofeedback, in [15] the feasibility of an online closed-loop system for AAD has been shown in an anechoic condition. Instead of using the clean speech signals of the attended and the unattended speaker as reference signals for decoding, in [16]–[20] the effect of different reference signals on the AAD performance has been investigated for an anechoic condition. Using simulated noisy reference signals for decoding, in [16] we have investigated the robustness of AAD to residual interference and background noise. In [17], [18] a neuro-steered noise reduction algorithm has been proposed to suppress the unattended speaker based on the AAD decision for an anechoic condition. In [19] an AAD-based sound source separation algorithm using deep neural networks has been presented to suppress the unattended speaker. In [20] we have investigated steerable beamformers to generate reference signals for AAD in an anechoic condition.

While the performance of the aforementioned least-squares-based AAD method has been extensively investigated for noiseless and anechoic acoustic conditions, in practice also background noise and reverberation, i.e. acoustic reflections against walls and objects, are present. Reverberation is known to spectro-temporally distort speech signals, causing the binaural spatial cues and pitch to become less reliable for performing auditory attention tasks [23]–[26]. In addition, interfering speakers and background noise degrade the attended speech signal, possibly leading to a severe speech encoding degradation at the level of the auditory nerve and the brainstem [27], [28]. Since in noisy and reverberant conditions the available signals at the ears contain several acoustic components (i.e. reverberation, background noise and interfering speaker), fully understanding the impact of each acoustic component on AAD is of crucial importance, e.g., in order to generate appropriate reference signals for decoding from these signals.

Recently, in [29] the performance of the least-squares-based AAD method was investigated for noisy and reverberant acoustic conditions. In [29] the same acoustic condition was used for AAD filter training and for decoding and the feasibility of using reverberant speech signals both as training and as reference signals was investigated. It was shown that in this way a comparable decoding performance for the reverberant condition as for the anechoic condition can be obtained. In this paper, we perform a more detailed analysis of the performance of the least-squares-based AAD method for an acoustic scenario comprising two competing speakers, background noise and reverberation. Compared to [29] we consider more acoustic conditions, especially with regard to background noise, and we specifically investigate the impact of different acoustic conditions for the training and the decoding steps. In addition, we investigate the influence on the decoding performance of the different acoustic components in the reference signals used for decoding and the training signals used for computing the filters. Some preliminary results were presented in [30], where we investigated the feasibility of using the (unprocessed) signals at the ears, containing reverberation, background noise and the interfering speaker, as reference and training signals.

The paper is organized as follows. In Section II the different acoustic conditions used for recording the EEG responses and the different acoustic signals used for the experimental analysis are introduced. In Section III the training and decoding steps of the least-squares-based AAD method are briefly reviewed. Section IV describes the acoustic and EEG measurement setup used for the experiments. In Section V the experimental results are presented and discussed, exploring the influence on the decoding performance of the different acoustic conditions and acoustic components.

## II. ACOUSTIC CONDITIONS AND COMPONENTS

We consider an acoustic scenario comprising two competing speakers and background noise in a reverberant environment (see Fig. 1). The clean speech signal of the attended speaker is denoted as *s^a^*[*i*], while the clean speech signal of the unattended speaker is denoted as *s^u^*[*i*], with *i* the discrete time index. The signals at the ears of the listener consist of a mixture of both speakers, including head filtering effects, reverberation and background noise. The signal *y_m_*[*i*] at the *m*-th ear, with *m* = 1 denoting the left ear and *m*=2 denoting the right ear, can be written as

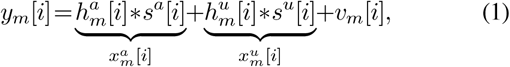

where 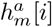 and 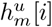 denote the (reverberant) acoustic impulse response between the *m*-th ear and the attended and the unattended speaker, respectively, * denotes the convolution operation, and *v_m_*[*i*] denotes the background noise component at the *m*-th ear. The reverberant speech signal of the attended and the unattended speaker at the *m*-th ear is denoted as 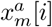 and 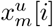, respectively. These reverberant speech signals consist of an anechoic speech signal encompassing the (anechoic) head filtering effect, i.e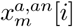 and 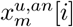, and a reverberation component. For notational conciseness the index *i* will be omitted in the remainder of this paper.

**Fig 1:**
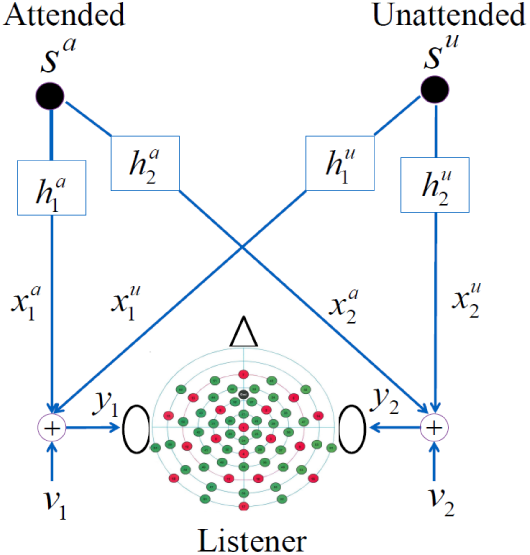
Binaural acoustic simulation setup used for simulating the presented stimuli in different acoustic conditions.

For the EEG recordings we will consider four different acoustic conditions, i.e. anechoic, reverberant, noisy and reverberant-noisy. Depending on the acoustic condition, the stimuli presented at the ears of the listener obviously comprise different acoustic components:

- in the *anechoic* condition (*an*), the mixture of the anechoic speech signals of the attended and the unattended speaker is presented.
- in the *noisy* condition (*no*), the mixture of the anechoic speech signals of the attended and the unattended speaker and background noise is presented.
- in the *reverberant* condition (*re*), the mixture of the reverberant speech signals of the attended and the unattended speaker is presented.
- in the *reverberant-noisy* condition (*rn*), the mixture of the reverberant speech signals of the attended and the unattended speaker and background noise is presented.

To investigate the impact of the different acoustic components on the AAD performance, we will consider several acoustic signals (see Table I):

- the clean speech signals *s^a^* and *s^u^*.
- the anechoic speech signals 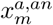and 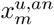, i.e. the clean speech signals affected by head filtering effects.
- the reverberant speech signals 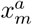 and 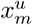, i.e. the anechoic speech signals affected by reverberation.
- the interfered speech signals, i.e. the anechoic speech signals affected by an interfering speaker

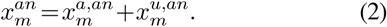
- the noisy speech signals, i.e. the anechoic speech signals affected by background noise

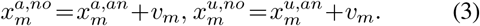
- the binaural speech signals *y_m_* in (1), i.e. the anechoic speech signals affected by reverberation, background noise and an interfering speaker.

**TABLE I:**
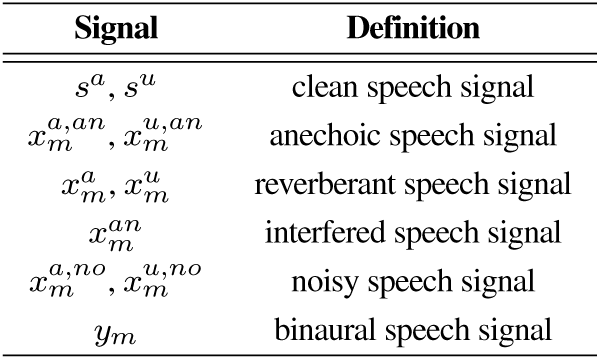
Acoustic Signals Used for Experimental Analysis

It should be noted that in the experiments (see Section IV-B) the positions of the attended and the unattended speaker are not always the same, i.e. for some participants the attended speaker is on the right side (and the unattended speaker on the left side), whereas for some participants the attended speaker is on the left side (and the unattended speaker on the right side). Due to the head filtering effect, the broadband energy ratio between the attended speech component and the unattended speech component in the signals at the ears is always smaller at the side of the unattended speaker than at the side of the attended speaker for the considered scenario. Therefore, the speech signals in Table I at the side of the attended speaker will be referred to as attended speech signals and the speech signals at the side of the unattended speaker as unattended speech signals.

## III. AUDITORY ATTENTION DECODING METHOD

This section briefly reviews the least-squares-based AAD method proposed in [10]. This method aims at reconstructing the attended speech envelope from the EEG recordings using a trained spatio-temporal filter. Section III-A describes the training step, where the envelope of a training signal is used together with the EEG recordings to compute the filter. Section III-B describes the decoding step, where the envelopes of two reference signals (attended and unattended) are compared with an estimate of the attended speech envelope computed using the trained filter. In most previous work [10]–[18], [21], [22] anechoic EEG recordings have been used in the training and decoding steps and/or the clean (or anechoic) speech signals have been used as training and reference signals. In this paper we will use EEG recordings from different acoustic conditions (see Section II) and we will explore the influence of using different acoustic signals (see Table I) as training and reference signals.

### A. Training Step

In the training step, the attended speaker is assumed to be known and an attended speech signal (e.g., the clean speech signal of the attended speaker *s^a^*) is used as training signal. From this signal the attended speech envelope *e^a^*[*k*], with *k* = 1…*K* the sub-sampled time index, is extracted, e.g., based on the Hilbert transform [31]. The attended speech envelope is then estimated from the EEG recordings *r_c_*[*k*], *c* = 1…*C*, using a spatio-temporal filter as

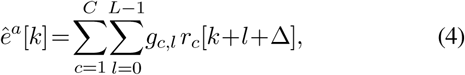

with *g_c,l_* the *l*-th filter coefficient in the *c*-th channel, *L* the number of filter coefficients per channel, and ∆ modeling the latency of the attentional effect in the EEG responses to the speech stimuli. In vector notation, (4) can be written as

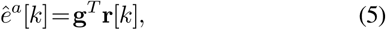

with

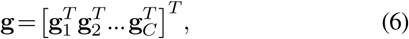

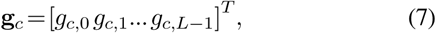

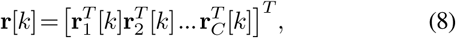

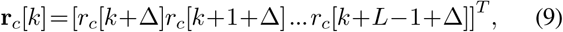

with (.)^*T*^ denoting the transpose operation. The spatio-temporal filter g is computed by minimizing the least-squares error between the attended speech envelope *e^a^*[*k*] and the reconstructed envelope *ê^a^*[*k*], regularized with the squared *l*_2_−norm of the derivatives of the filter coefficients to avoid over-fitting [12], [30], i.e.

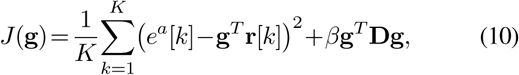

with **D** denoting the derivative matrix and *β* denoting a regularization parameter. The filter minimizing the regularized least-squares cost function in (10) is equal to

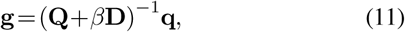

with the correlation matrix **Q** and the cross-correlation vector **q** given by

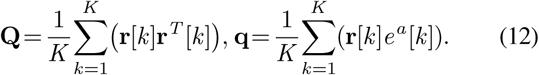

In this paper we will consider several training conditions (*tc*) for computing the filter g, i.e. *tc* = *an* using EEG responses recorded in the anechoic condition, *tc* = *re* using EEG responses recorded in the reverberant condition, *tc* = *no* using EEG responses recorded in the noisy condition, and *tc* = *rn* using EEG responses recorded in the reverberant-noisy condition. In addition, we will consider the training condition *tc* = *ac*, in which EEG responses from all conditions are used for computing the filter.

Aiming at investigating the influence of each acoustic component, in this paper we will consider different attended speech signals (see Table I) as training signals, more in particular the clean attended speech signal *s^a^*, the anechoic attended speech signal 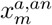 the reverberant attended speech signal 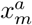, the interfered attended speech signal 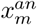 the noisy attended speech signal 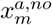 and the binaural attended speech signal *y_m_*.

### B. Decoding Step

For each acoustic condition, the complete set of EEG responses is segmented into *T* trials (see Section IV-C for more details). The filter corresponding to trial *t* to be decoded is denoted as g*_t_*. To decode to which speaker a listener attended during trial *t*, first an estimate of the attended speech envelope 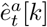 is computed using the (trained) filter g*_t_*, i.e.

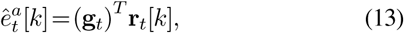

with **r***_t_* [*k*] denoting the EEG recordings of trial *t*. Next, the correlation coefficients between the estimated attended speech envelope 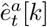 and the envelope of two reference signals, i.e. namely the attended and the unattended reference signal, are computed as

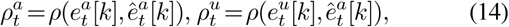

where 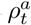and 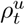 denote the attended and the unattended correlation coefficient, respectively, and 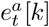 and 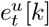 denote the attended and the unattended speech envelope, respectively. When 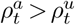, it is decided that auditory attention has been correctly decoded. Accordingly, a larger difference between the attended and the unattended correlation coefficient 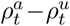 (referred to as correlation difference) is indicative of a more reliable AAD decision. The decoding performance *P* is defined as the percentage of correctly decoded trials over all considered trials and all participants. To compute the correlation coefficients in (14), EEG recordings in different acoustic conditions can be used for computing 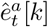In addition, aiming at investigating the influence of each acoustic component on the decoding performance, different reference signals (see Table I) can be used for computing the attended and the unattended speech envelope 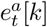 and 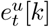, respectively.

In this paper we will investigate the decoding performance for several evaluation conditions *ec* ∈ {*an,re,no,rn,ac*}, with *P_ec_* denoting the decoding performance for a specific evaluation condition. To decode trial *t* of an evaluation condition using the filter trained in a specific training condition which is not necessary the same as the evaluation condition, the filter g*_t_* is computed as follows:

- when the trial *t* to be decoded is part of the trials in the training condition, the filter is computed using (11) as

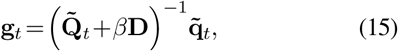

with 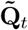 the average correlation matrix, computed by averaging all correlation matrices corresponding to trials in the training condition *except* trial *t*, and 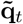 the average cross-correlation vector, computed by averaging all cross-correlation vectors corresponding to trials in the training condition *except* trial *t*, i.e.

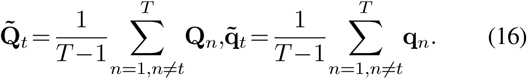

This procedure corresponds to leave-one-out averaging.
- when the trial *t* to be decoded is not part of the trials in the training condition, the filter is computed using (11) as

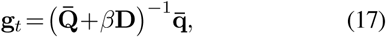

with 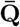 the average correlation matrix, computed by averaging all correlation matrices corresponding to trials in the training condition, and 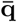 the average cross-correlation vector, computed by averaging all cross-correlation vectors corresponding to trials in the training condition, i.e.

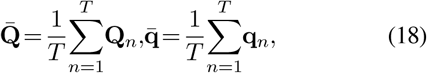

Since the number of trials across acoustic conditions is different (see Section IV-B), for *tc* = *ac* the average correlation matrix and the average cross-correlation vector (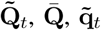 and 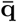) are computed in such a way that the contribution of trials from each acoustic condition is considered equally.

In [16] it has been shown that the parameters involved in the filter design (∆, *L*, *β*) play an important role in obtaining a good decoding performance. In order not to favour one specific acoustic evaluation condition, the filter parameters have been determined to optimize the average decoding performance *P_ac_* over all considered acoustic conditions. Please note that the filter parameters have been optimized per participant and for each training condition (see Section IV-B and IV-C).

## IV. ACOUSTIC AND EEG MEASUREMENT SETUP

### A. Participants

Eighteen native German-speaking participants (right-handed and aged between 21 and 34 years) took part in this study. All participants were normal-hearing as was confirmed by pure tone audiometry. The participants reported no past or present neurological or psychiatric conditions. All participants signed an informed consent form and were paid for their participation. Two participants were excluded from the analysis, one participant due to poor attentional performance (as revealed by the questionnaire results) and the other participant due to a technical hardware problem.

### B. Acoustic Stimuli

Two German audio stories, uttered by two different male speakers, were used as the clean speech signals (sampling frequency of 16 kHz). One story was from the German audio book website [35] and the other story was from a selection of audio books [36]. Speech pauses that exceeded 0.5 s were shortened to 0.5 s. The acoustic stimuli were simulated by convolving the clean speech signals (i.e. the audio stories) with non-individualized binaural acoustic impulse responses, either from [32], [33], or [34], and by adding diffuse babble noise, generated according to [37]. The competing speakers were simulated at −45° (left) and 45° (right). Eight different acoustic conditions were considered for the stimuli (see Table II): anechoic, reverberant with a moderate and a large reverberation time (*T*_60_ = 0.5 s, *T*_60_ = 1 s), noisy with two different broadband signal-to-noise ratios (SNR = 9.0 dB, SNR = 4.0 dB), and three combinations of reverberation and noise. For the experimental analysis, the acoustic conditions were grouped based on acoustic similarity as shown in Table II, resulting in four experimental analysis conditions, i.e. anechoic, reverberant, noisy, and reverberant-noisy. The acoustic stimuli were presented to the participants via insert earphones (E-A-RTONE 3A) using an RME HDSP 9632 PCI Audio Interface and Tucker Davis Technologies programmable attenuators.

**TABLE II:**
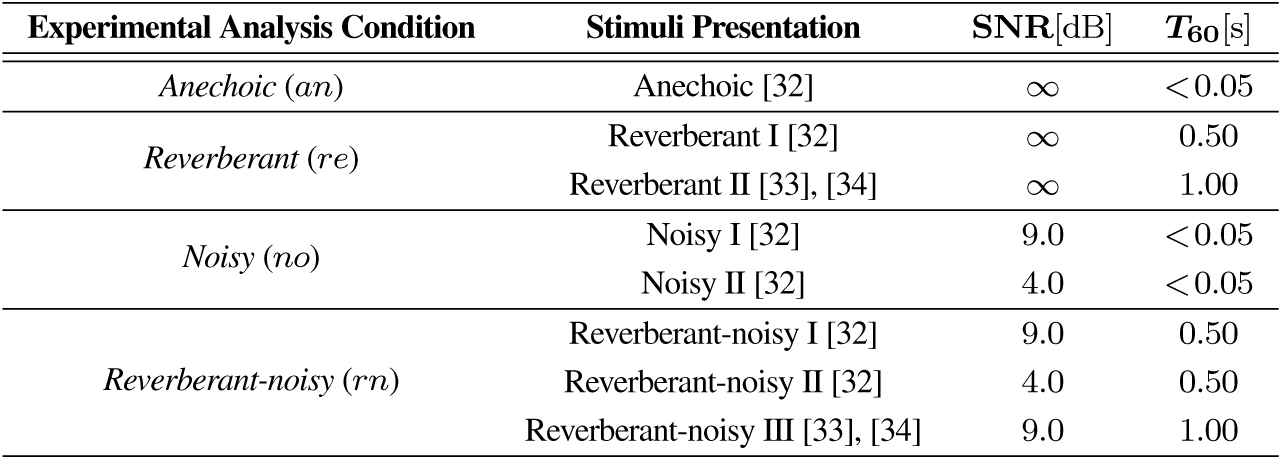
Acoustic conditions used for experimental analysis and stimuli presentation

Before performing the experiment, the participants reported no, or very limited, knowledge of the audio stories. Among all participants, 8 participants were instructed to attend to the left speaker, while 10 participants were instructed to attend to the right speaker. The participants were instructed to look at a fixation cross on a screen and minimize eye blinking. The stimuli were presented in 11 sessions, each of length 10 minutes, interrupted by short breaks. For each participant, the anechoic condition was always assigned to the first session and subsequently to every other third session (i.e. session 4, 7, and 10). Aiming at minimizing the influence of the speech material on AAD, the acoustic conditions (except for the anechoic condition) were randomly assigned to the other sessions. Following each session, the participants were asked to fill out a questionnaire consisting of 10 multiple-choice questions related to each story. The questionnaire was aimed to indicate whether the participants attended to the instructed speaker and whether the audio story was intelligible in the different acoustic conditions. The experiment for each participant took place on two different days.

### C. EEG Setup and Signal Pre-processing

The EEG responses were recorded using *C* = 64 channels, provided by Easycap GmbH, Germany, with a sampling frequency of 500 Hz. The EEG responses were referenced to the nose electrode and recorded using the Brain-Vision recorder software. The EEG recordings were re-referenced offline to a common average reference, band-pass filtered between 2 Hz and 8 Hz using a third-order Butterworth band-pass filter, and subsequently downsampled to *f_s_* = 64 Hz. The envelopes of all considered 16 kHz speech signals were obtained using a Hilbert transform [31], followed by low-pass filtering at 8 Hz and downsampling to *f_s_* = 64 Hz. For the training and decoding steps (see Section III), the EEG recordings of each session were split into 10 trials, each of length 60 seconds. For filter training and evaluation, each participant’s own data were used.

## V. RESULTS AND DISCUSSION

In this section, the decoding performance of the least-squares-based AAD method is investigated for different acoustic conditions (see Table II) using the experimental setup discussed in the previous section. Section V-A discusses the results of the questionnaire. In Section V-B the impact of different acoustic conditions for the training and decoding steps is investigated. In Section V-C the impact of the head filtering effect is explored by comparing the decoding performance using either the clean or the anechoic speech signals. Finally, in Section V-D the influence of each acoustic component is investigated by comparing the decoding performance using reference and training signals affected by background noise, reverberation, and/or interfering speaker.

### A. Questionnaire Analysis

For all considered acoustic conditions, Fig. 2 presents the correct answer scores related to the attended story, averaged across all participants. The highest score is obtained for the anechoic condition, while the lowest score is obtained for the reverberant-noisy condition. The statistical multiple comparison test (Kruskal-Wallis test followed by post-hoc Dunn and Sidak test [38]) showed a significant difference (Kruskal-Wallis test: χ^2^ = 19.0, *p* = 0.002) in terms of the correct answer score between the anechoic condition and either the noisy or the reverberant-noisy condition (post-hoc Dunn and Sidak test: *p*=0.022 and *p* = 0.000, respectively) and between the reverberant condition and the reverberant-noisy condition (post-hoc Dunn and Sidak test: *p* = 0.013), implying that – as expected – the noisy and the reverberant-noisy condition are more challenging.

**Fig 2:**
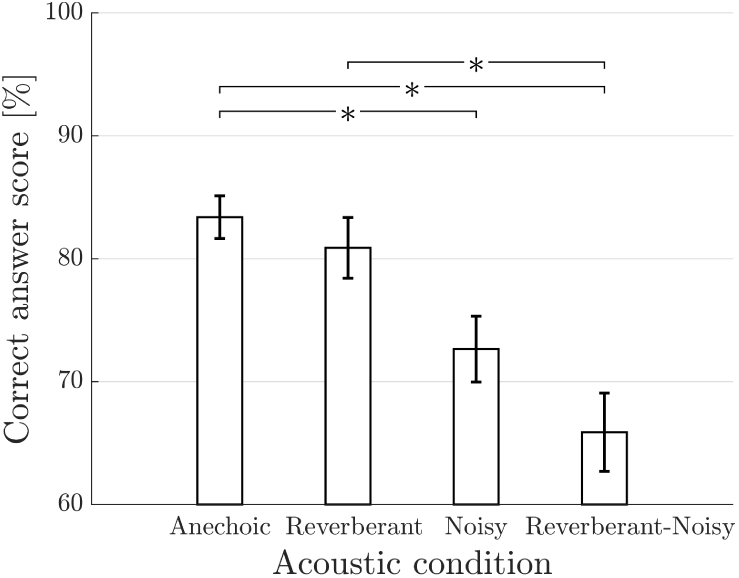
The correct answer scores related to the attended story, averaged across all participants, for different acoustic conditions. Error bars represent one standard error around the mean and * indicates a significant difference (*p* < 0.05) between acoustic conditions, based on the Kruskal-Wallis test followed by the post-hoc Dunn and Sidak test.

### B. Impact of Acoustic Conditions

For all considered evaluation conditions, Fig. 3 presents the decoding performance for different training conditions when the clean speech signals are used as reference and training signals.

**Fig 3:**
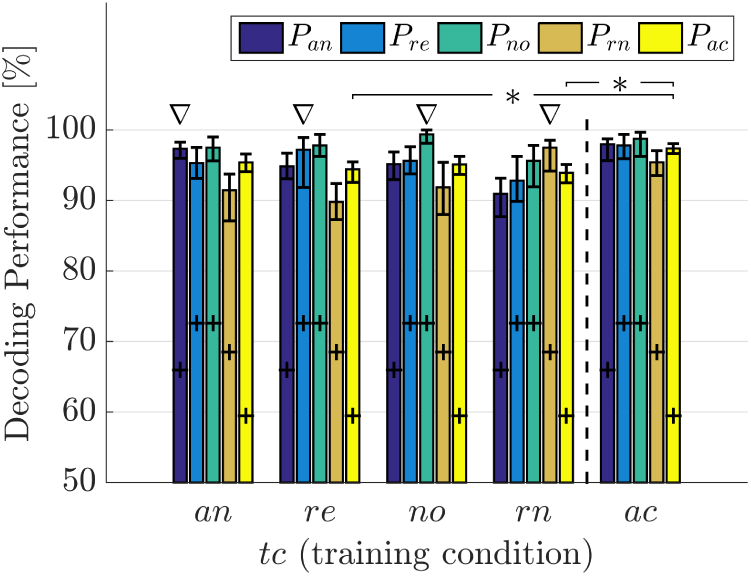
The decoding performance for different training and evaluation conditions when using the clean speech signals. The plus signs represent the upper boundary of the confidence interval corresponding to chance level, based on a binomial test at the 5% significance level, the error bars represent the bootstrap confidence interval at the 5% significance level, ∇ indicates the decoding performance when the training and evaluation conditions are equal, and * indicates a significant difference (*p* < 0.05) with the decoding performance for all conditions (*P_ac_*) based on the Kruskal-Wallis test followed by the post-hoc Dunn and Sidak test.

First, we investigate the feasibility of decoding EEG responses in different acoustic conditions *ec*∈{*an,re,no,rn,ac*} when using filters trained using EEG responses in a *specific* acoustic condition *tc* ∈ {*an,re,no,rn*} (i.e. left part of Fig. 3, separated by dashed line). When the evaluation and training conditions are equal (indicated by ∇), it can be observed that a very good decoding performance (> 96%) is obtained for all evaluation conditions. These results are consistent with previous findings for the anechoic condition [11], [13],[15]–[18] as well as with recent findings for the reverberant and the reverberant-noisy conditions [30]. For each training condition *tc* ∈ {*an,re,no,rn*}, it can be observed that the decoding performance when the evaluation and training conditions are equal (indicated by ∇) is among the highest decoding performances for all evaluation conditions. When the evaluation and training conditions are not equal, typically a lower decoding performance is obtained (except in some cases for the anechoic and the reverberant training conditions). For example, for the reverberant-noisy training condition the highest decoding performance is obtained for the reverberant-noisy evaluation condition (> 97%), while a lower decoding performance is obtained for the anechoic, reverberant, and noisy evaluation conditions (> 90%). In addition, for all training conditions *tc* ∈{*an,re,no,rn*} it can be observed that the average decoding performance for all conditions *P_ac_* is considerably high (> 93%).

Secondly, we investigate the feasibility of decoding EEG responses in different acoustic conditions *ec* ∈ {*an,re,no,rn,ac*} when using filters trained using EEG responses in *all acoustic conditions tc* = *ac* (i.e. right part of Fig. 3, separated by dashed line). It can be observed that a very good decoding performance (> 95%) is obtained for all evaluation conditions and that the decoding performance across evaluation conditions is more consistent compared to when using filters trained in a specific acoustic condition. In addition, the average decoding performance for all conditions *P_ac_* obtained with filters trained in all conditions is occasionally significantly larger than with filters trained in a specific acoustic condition. For example, the decoding performance *P_ac_* obtained with filters trained in all conditions (*tc* = *ac*) is significantly larger than with filters trained either in the reverberant condition (*tc* = *re*) or in the reverberant-noisy condition (*tc* = *rn*) (Kruskal-Wallis test: χ^2^ =16.5, *p* = 0.002; post-hoc Dunn and Sidak test comparisons of *tc* = *ac* with *tc* = *re* and *tc* = *rn*: *p* = 0.020, *p* = 0.001, respectively).

The feasibility of using either filters trained in a specific acoustic condition or filters trained in all acoustic conditions to perform AAD in different acoustic conditions may be explained by considering the robust neural responses to degraded – but still intelligible – speech signals. Several studies have shown that auditory cortical responses resemble the clean attended speech signal more than the speech signal degraded by different acoustic components (e.g., background noise, interfering speaker), suggesting a robust neural representation of the clean attended speech signal [6], [7], [28], [29], [39]. To decode auditory attention, the trained filters aim at reconstructing the clean attended speech envelope from EEG responses that are largely invariant to degradations. Hence, the reconstructed attended envelope is expected to be more correlated to the clean attended speech envelope than to the clean unattended speech envelope, i.e. the correlation difference (*ρ^a^*−*ρ^u^*) is expected to be larger than zero. For all considered evaluation conditions, Fig. 4 presents the correlation difference for different training conditions, averaged across all considered trials and participants (note that these average correlation coefficients are not directly used for decoding). It can be observed that a correlation difference significantly larger than zero is obtained for all considered acoustic conditions, which is consistent with a robust neural representation of the clean attended speech signal.

**Fig 4:**
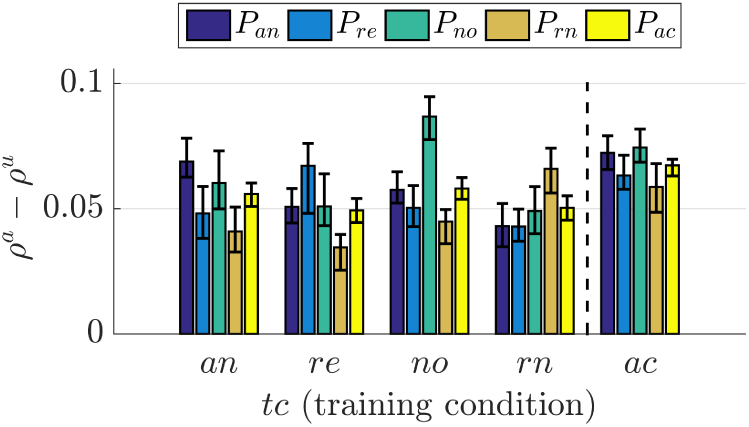
Average correlation differences for different training conditions and evaluation conditions when using the clean speech signals. The error bars represent the bootstrap confidence interval at the 5% significance level.

Finally, we investigate the parameters involved in the filter design (∆, *L*, *β*) across training conditions. Fig. 5 depicts the optimal parameter values (see Section III-B), averaged across all considered trials and all participants. It can be observed that the optimal value for ∆ varies only slightly between 93.8 ms to 101.6 ms, while the optimal value for ∆ varies more substantially between 109.3 ms to 128.9 ms. Accordingly, the EEG responses contributing most to the AAD performance are those with latencies between 93.8 ms and 230.5 ms, consistent with previous findings in [12], [16], [28]. In addition, the optimal value for the regularization parameter varies between 10^−1^ to 10^2^. It can be observed that the optimal regularization parameter is smaller when using filters trained in all conditions than when using filters trained in a specific acoustic condition. A possible explanation may be that training in all conditions can by itself be considered as some form of regularization.

**Fig 5:**
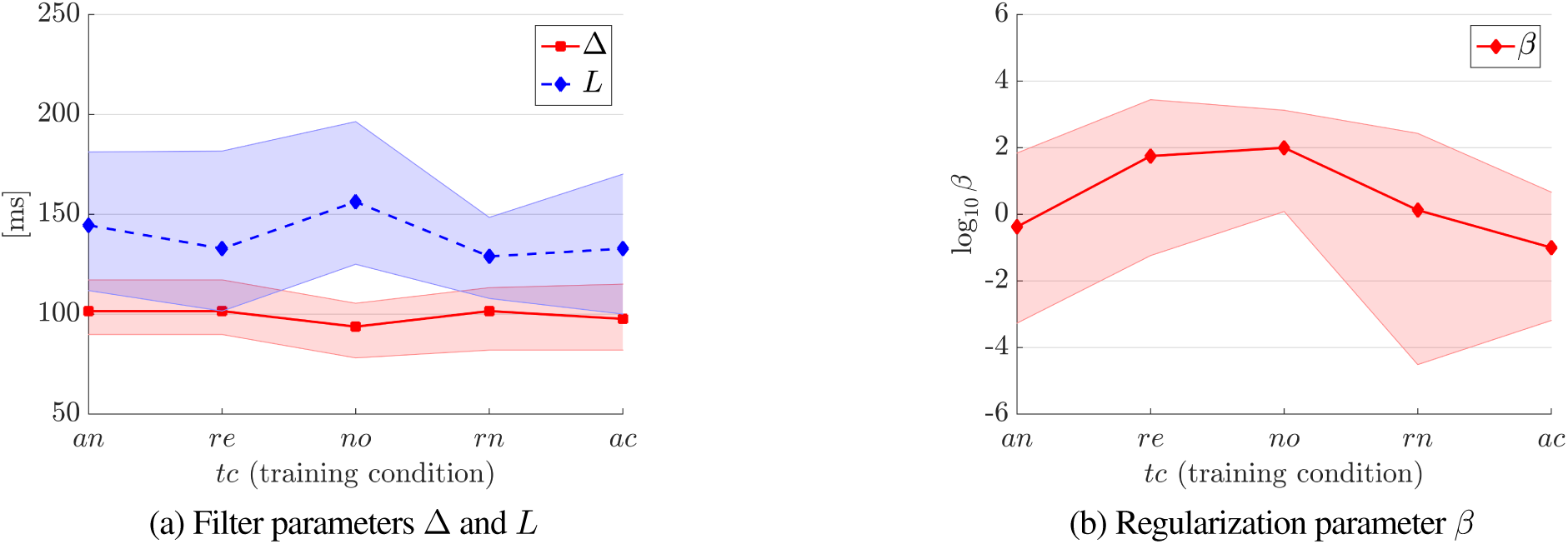
The optimal values for the filter parameters (a) ∆ and *L*, and (b) the regularization parameter *β*, averaged across all trials and all participants when using the clean speech signal. The shaded area indicates the bootstrap confidence interval at the 5% significance level.

In summary, the results in this section show the feasibility of using either filters trained in a specific acoustic condition or filters trained in all conditions to perform AAD in different acoustic conditions. While these results were obtained using the clean speech signals as training and reference signals, in the next sections we will investigate in more detail the influence of the different acoustic components (head filtering effect, reverberation, background noise, interfering speaker) in the training and reference signals.

### C. Influence of head filtering effect

In this section, we investigate the influence of the head filtering effect by comparing the decoding performance when using clean or anechoic speech signals either as training or as reference signals. Fig. 6a presents the decoding performance for the anechoic condition (*ec* = *an*) when using filters trained in the anechoic condition (*tc* = *an*). Fig. 6b presents the average decoding performance for all conditions (*ec* = *ac*) when using filters trained in all conditions (*tc* = *ac*). A paired Wilcoxon signed rank test revealed no significant difference (*p* > 0.05) between using either the clean speech signals or the anechoic speech signals as training or as reference signals. These results indicate that for all considered acoustic conditions head filtering effects have no significant influence on the decoding performance.

**Fig 6:**
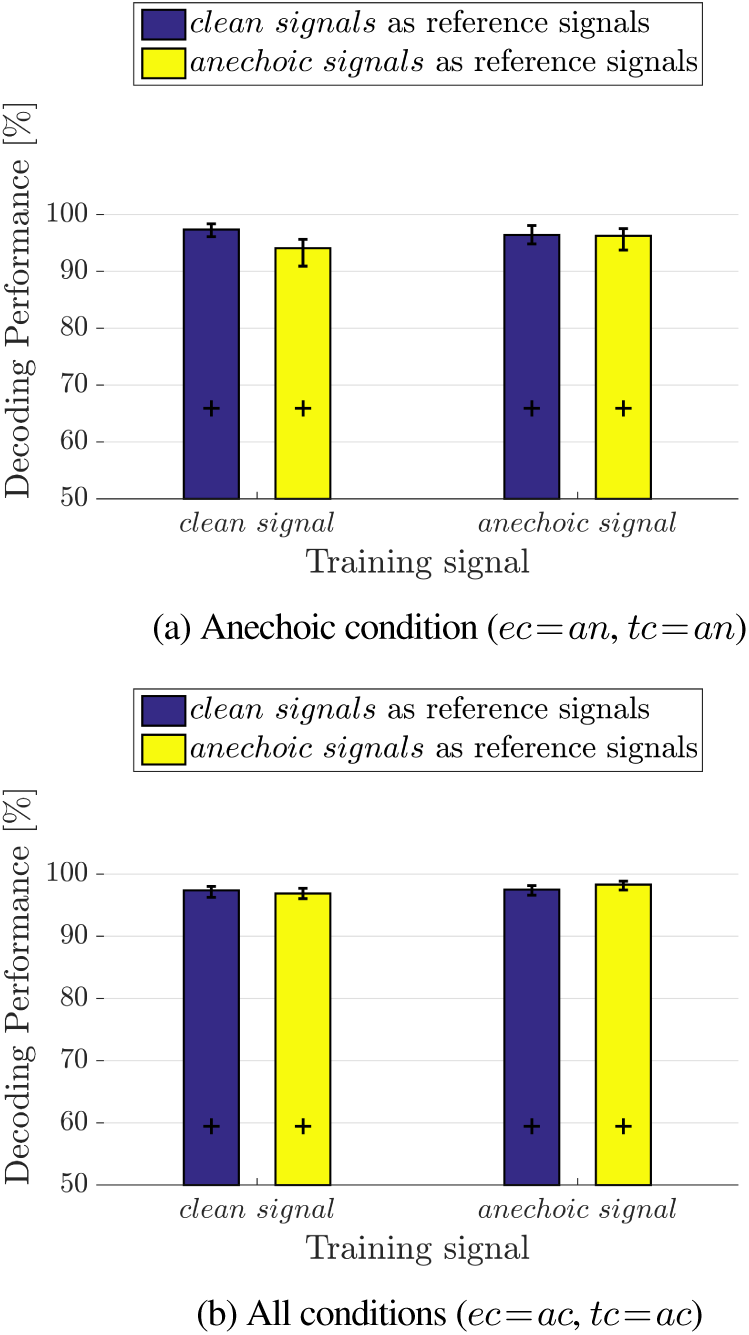
Influence of head filtering effect on AAD. Comparison of decoding performance using either the clean or the anechoic speech signals when the evaluation and training conditions are equal to (a) the anechoic condition or (b) all conditions. The plus signs represent the upper boundary of the confidence interval corresponding to chance level based on a binomial test at the 5% significance level, the error bars represent the bootstrap confidence interval at the 5% significance level.

### D. Influence of background noise, reverberation and interfering speaker

To investigate the influence of each acoustic component on AAD, Fig. 7 presents the decoding performance for all considered acoustic conditions (anechoic, reverberant, noisy, reverberant-noisy) using the following signals as training signals or as reference signals:

- the clean speech signals *s^a^* and *s^u^*.
- the anechoic speech signals 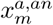 and 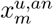.
- the anechoic speech signals affected by different acoustic components, i.e. the noisy speech signals 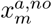and 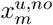 in (3) for the noisy condition, the reverberant speech signals 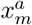 and 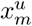 in (1) for the reverberant condition, the interfered speech signal 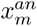 (attended and unattended side) in (2) for the anechoic condition^1^, and the binaural speech signals *y_m_* (attended and unattended side) in (1) for the reverberant-noisy condition.

**Fig 7:**
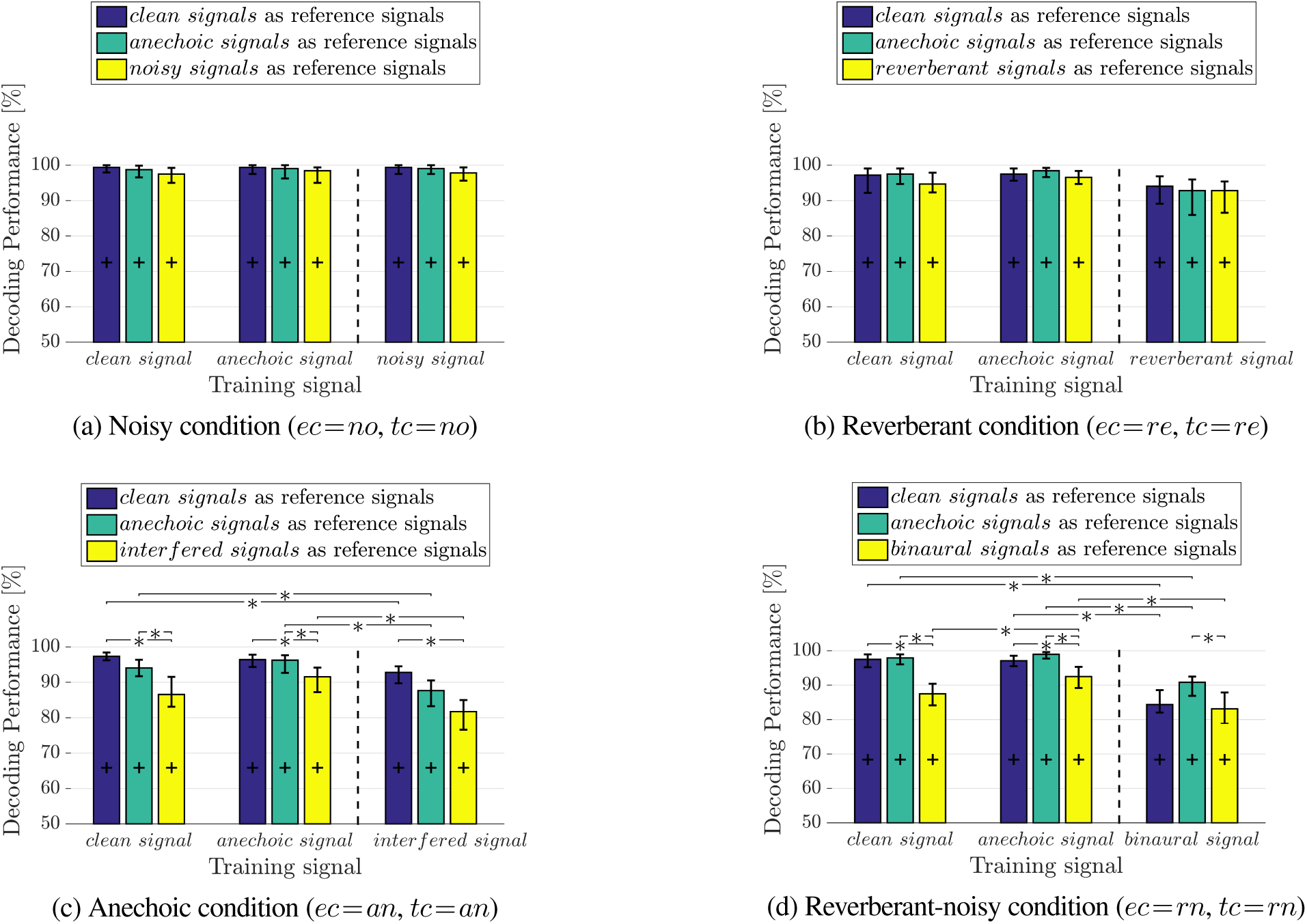
Influence of different acoustic components (background noise, reverberation and interfering speaker) on AAD. Comparison of decoding performance when using (a) the noisy speech signals in the noisy condition, (b) the reverberant speech signals in the reverberant condition, (c) the interfered speech signals in the anechoic condition, (d) the binaural speech signals in the reverberant-noisy condition, either as training signal or as reference signals. The plus signs represent the upper boundary of the confidence interval corresponding to chance level based on a binomial test at the 5% significance level, the error bars represent the bootstrap confidence interval at the 5% significance level, and indicates a significant difference (*p* < 0.05) based on the paired Wilcoxon signed rank test.

Similarly, Fig. 8 presents the correlation difference (*ρ^a^* − *ρ^u^*), averaged across all considered trials and participants (note that these average correlation coefficients are not directly used for decoding).

**Fig 8:**
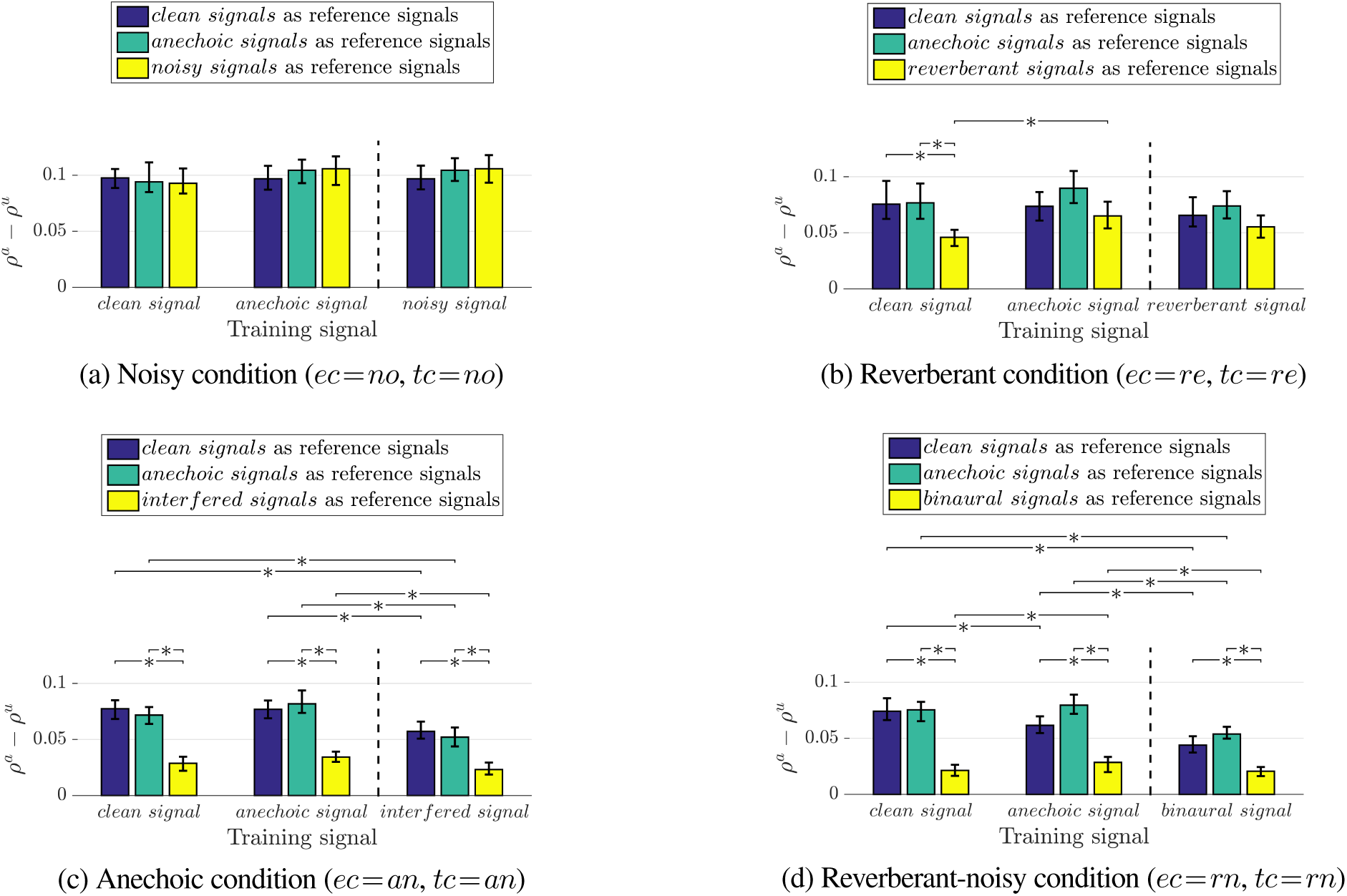
Influence of different acoustic components (background noise, reverberation and interfering speaker) on AAD. Comparison of correlation difference when using (a) the noisy speech signals in the noisy condition, (b) the reverberant speech signals in the reverberant condition, (c) the interfered speech signals in the anechoic condition, (d) the binaural speech signals in the reverberant-noisy condition, either as training signal or as reference signals. The error bars represent the bootstrap confidence interval at the 5% significance level, and indicates a significant difference (*p* < 0.05) based on the paired Wilcoxon signed rank test.

First, we investigate the case where the clean or the anechoic attended speech signal is used as training signal (i.e. left part of Fig. 7 and 8, separated by dashed line). When using the clean or anechoic speech signals as reference signals, a very good decoding performance (> 94%) is obtained for all acoustic conditions, as already shown in Fig. 3. When using the noisy speech signals (in the noisy condition, Fig. 7a) or the reverberant speech signals (in the reverberant condition, Fig. 7b) as reference signals, there is no significant difference in decoding performance (*p* > 0.05) compared to when using the clean or anechoic speech signals as reference signals. On the other hand, when using the interfered speech signals (in the anechoic condition, Fig. 7c) or the binaural speech signals (in the reverberant-noisy condition, Fig. 7d) as reference signals, the decoding performance is significantly lower (*p* < 0.05) than when using the clean or anechoic speech signals as reference signals, although the decoding performance is still considerably large (> 87%). The feasibility of using either the interfered speech signals or the binaural speech signals as reference signals for AAD can be explained by considering the broadband energy ratio between the attended and unattended speech components in the signals at the ears. As already mentioned in Section II, due to the head filtering effect this broadband energy ratio is smaller at the side of the unattended speaker than at the side of the attended speaker. In summary, the results in Fig. 7 (left side) show that when using reference signals affected by reverberation or background noise, a comparable decoding performance can be obtained as when using clean or anechoic speech signals, whereas when using reference signals affected by the interfering speaker the decoding performance significantly decreases. This also suggests that in order to generate appropriate reference signals, it is more important to reduce the interfering speaker than to reduce background noise or reverberation.

The decoding performance results in Fig. 7 can be further explained by considering the influence of each acoustic component on the correlation difference in Fig. 8. For the noisy condition (Fig. 8a), there are no significant differences between the considered reference signals, which corresponds to the decoding performance results in Fig. 7a. For the reverberant condition (Fig. 8b), it can be observed that the correlation differences significantly decrease (*ρ^a^* − *ρ^u^* < 0.04) when using the reverberant speech signals as reference signals, but only when using the clean attended speech signal as training signal. Nevertheless, this lower correlation difference does not result in a significantly lower decoding performance in Fig. 7b. For the anechoic condition (Fig. 8c) and the reverberant-noisy condition (Fig. 8d), it can be observed that the correlation differences significantly decrease when using the interfered speech signals (*ρ^a^* − *ρ^u^* <0.03) or the binaural speech signals (*ρ^a^* − *ρ^u^* < 0.02) as reference signals. These lower correlation differences are also reflected by significantly lower corresponding decoding performances in Fig. 7c and 7d.

Secondly, we explore the potential of using the attended speech signal affected by different acoustic components as training signal (i.e. right part of Fig. 7 and 8, separated by dashed line). On the one hand, when using the noisy attended speech signal (in the noisy condition, Fig. 7a) or the reverberant attended speech signal (in the reverberant condition, Fig. 7b) as training signal, there is no significant difference in decoding performance (*p* > 0.05) compared to when using the clean or the anechoic attended speech signal as training signal (for all considered reference signals). On the other hand, when using the interfered attended speech signal (in the anechoic condition, Fig. 7c) or the binaural attended speech signal (in the reverberant-noisy condition, Fig. 7d) as training signal, the decoding performance is significantly lower compared to when using either the clean or the anechoic attended speech signal as training signal (for all considered reference signals). Nevertheless, even when using the binaural attended speech signal as training signal in the reverberant-noisy condition, it is still feasible to perform AAD with a decoding performance larger than 82%. The decoding performance results in Fig. 7 when using attended speech signals affected by different acoustic components as training signal are mostly consistent with the correlation differences in Fig. 8.

In summary, the results in this section show that using speech signals affected by background noise and reverberation as training or reference signals results in a decoding performance that is comparable to using the clean or anechoic speech signals as training or reference signals. On the contrary, using speech signals affected by the interfering speaker as training or reference signals typically results in a significantly lower decoding performance.

## VI. CONCLUSIONS

In this paper, we investigated the performance of the least-squares-based AAD method for different acoustic conditions (anechoic, reverberant, noisy, and reverberant-noisy), both in the training step as well as in the decoding step. The experimental results showed that for all considered acoustic conditions it is possible to decode auditory attention with a decoding performance larger than 90%, even when the acoustic conditions for training and decoding are different. In addition, for most acoustic conditions there is no significant difference in decoding performance when using filters trained in all conditions or filters trained in a specific condition.

Furthermore, we investigated the influence of the head filtering effect and of acoustic components (reverberation, background noise and interfering speaker) on the decoding performance. The experimental results showed that for all considered acoustic conditions the head filtering effect has no significant impact on the decoding performance. Moreover, when using speech signals affected by either reverberation or background noise as reference signals, a comparable decoding performance is obtained as when using clean speech signals as reference signals. On the contrary, when using speech signals affected by the interfering speaker as reference signals, the decoding performance significantly decreases. Nevertheless, even when using the binaural speech signals as reference signals for decoding, a relatively large decoding performance can be obtained.

Finally, we explored the potential of using the attended speech signal affected by different acoustic components as training signal for computing the filter. When using attended speech signals affected by either reverberation or by background noise as training signal, a comparable decoding performance is obtained as when using the clean attended speech signal as training signal. However, when using attended speech signals affected by the interfering speaker as training signal, the decoding performance may significantly decrease. Nevertheless, even when using the binaural attended speech signal as training signal, it is still feasible to achieve a decoding performance larger than 82%.

While the discussion in this paper has been limited to the least-squares-based AAD method, in which auditory attention is decoded using an envelope reconstruction model, AAD approaches based on a neural encoding model [40] have not been investigated in this paper. Further work could therefore include a study on how reverberation and noise influence these neural-encoding-based AAD approaches.

## Footnotes

1 The interfered speech signal is used in the anechoic condition to exclude the influence of other acoustic components (background noise and reverberation) on the analysis.

